# Challenges in reconstituting the peroxiredoxin 2:STAT3 transient redox-relay complex *in vitro*

**DOI:** 10.64898/2026.07.03.731362

**Authors:** Julia Malo Pueyo, Ekaterina Baranova, Khadija Wahni, Victor R. A. Dubach, Steven Janvier, Didier Vertommen, Bonnie J. Murphy, Daria Ezeriņa, Joris Messens

## Abstract

Peroxiredoxin 2 (Prdx2) mediates redox signaling by transferring oxidative equivalents to target proteins such as STAT3, a redox-sensitive transcription factor implicated in inflammation and cancer. Although this interaction has been demonstrated in cells, reconstituting the Prdx2:STAT3 complex *in vitro* remains challenging due to its transient and redox-dependent nature. Here we test various conditions to stabilize the complex between taggless Prdx2 and the core fragment of STAT3 (CF-STAT3), including oxidants, detergents, the facilitator Annexin A2, anaerobic environments, and CovalX crosslinking. Complex formation was assessed via mass photometry, analytical size-exclusion chromatography (SEC), SEC-MALS, and electron microscopy (EM). No stable complex was observed under standard conditions. Anaerobic environments briefly stabilized the interaction, but cryo-EM could not resolve the structure. CovalX crosslinking yielded short-lived but homogeneous complexes. We found that Prdx2 is highly susceptible to hyperoxidation at its peroxidatic cysteine, particularly in the presence of DTT or excess H_2_O_2_, resulting in loss of function. Maintaining non-reducing conditions during purification preserved Prdx2 in an oxidation-competent state, promoting formation of the disulfide bond between the peroxidatic and resolving cysteines and thereby enabling reproducible detection of a weak complex with CF-STAT3. Our findings establish a framework for studying redox-relay protein complexes *in vitro* and highlight the importance of oxidation state management during protein handling.

## 1. Introduction

Peroxiredoxins (Prdxs) are thiol-based peroxidases that play a central role in detoxifying hydrogen peroxide (H_2_O_2_) and mediating redox signaling (Figure 1). Prdxs function through catalytic cycling of a conserved peroxidatic cysteine (Cys_P_) and a resolving cysteine (Cys_R_), which form an intermolecular disulfide bond ^1-4^. This catalytic cycle is then completed by the reduction of the intersubunit disulfide bond via the thioredoxin (Trx)–thioredoxin reductase (TrxR)-nicotinamide adenine dinucleotide phosphate (NADPH) system ^5,6^. At elevated H_2_O_2_ concentrations, the conserved peroxidatic cysteine (Cys_P_), already oxidized in the sulfenic form, can undergo further oxidation to sulfinic or sulfonic acid forms — collectively referred to as hyperoxidation. These modifications inactivate their peroxidase function, allowing H_2_O_2_ levels to rise and promoting oxidation of other cellular components, such as proteins, lipids, and DNA, contributing to oxidative stress. Notably, some hyperoxidized Prdxs gain chaperone-like activity, functioning as high-molecular-weight holdases under stress conditions ^4^. Prdxs not only regulate intracellular peroxide levels but also function as redox sensors and signaling mediators ^1^. Over the past decades, 2-Cys Prdxs have been shown to transfer oxidative equivalents to specific target proteins through redox-relay mechanisms, extending their role beyond peroxide detoxification ^7-11^. For instance, one target of Prdx2 is STAT3 (Signal Transducer and Activator of Transcription 3), a transcription factor central to inflammation, cell proliferation, and cancer ^12-14^. Oxidation of STAT3 by Prdx2 affects its dimerization and DNA-binding activity, thereby modulating its transcriptional output in response to oxidative signals ^10^. Annexin A2 (AnxA2) has been proposed to facilitate this redox relay by scaffolding or stabilizing the Prdx2:STAT3 interaction in cells ^10^. However, the structural and biochemical details of the Prdx2:STAT3 complex remain unclear. This study aims to outline the challenges of reconstituting the Prdx2:STAT3 complex *in vitro*.

**Figure 1.**
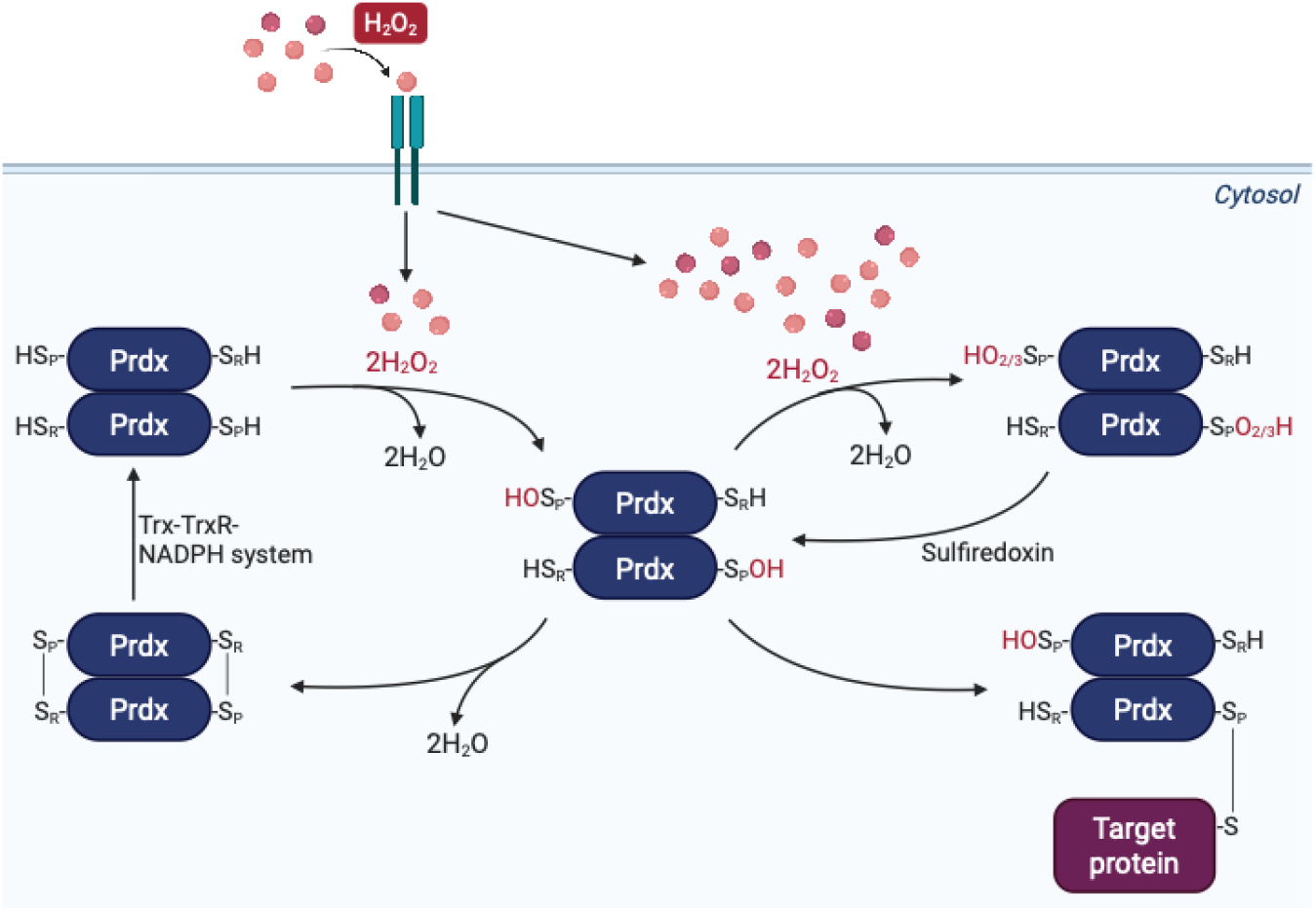
Typical 2-Cys Prdxs perform multiple functions inside cells. Reduced Cys_P_-SH from one subunit of the obligate homodimer can react with a H_2_O_2_ molecule, becoming oxidized to the sulfenic acid (Cys_P_-SOH) form. Then, the Cys_R_-SH of the other subunit of the obligate homodimer undergoes a condensation reaction with the Cys_P_-SOH to form an intersubunit disulfide bond (Cys_P_-S-S-Cys_R_), which will subsequently be reduced by the Trx-TrxR-NADPH system. On the other hand, at high H_2_O_2_ concentrations, the oxidized CysP-SOH residue can alternatively become hyperoxidized to the sulfinic or sulfonic form (Cys_P_-SO_2/3_). Oxidized peroxidatic Cys can also form a mixed disulfide intermediate with a target binding partner protein. Figure created using BioRender.

## 2. Materials and Methods

### 2.1. Antibodies, chemicals, and plasticware

Antibodies were rabbit anti-Prdx2, kindly provided by the lab of Bernard Knoops (Université Catholique de Louvain, Belgium) ^15^, mouse anti-His (AD1.1.10, MCA1396, Bio-Rad Laboratories, Richmond, VA, USA), anti-mouse IgG (A0168, Sigma-Aldrich, St. Louis, MO, USA) and anti-rabbit IgG (12-348, EMD Millipore Corp, USA). The protein ladder used for sodium dodecyl sulfate– polyacrylamide gel electrophoresis (SDS-PAGE) was PageRuler Prestained protein ladder (26616, Thermo Fisher Scientific, Waltham, WA, USA). All chemicals were purchased from Merck (Sigma-Aldrich, St. Louis, MO, USA) unless stated otherwise. Antibiotics and dithiothreitol (DTT) were purchased from Duchefa Biochemie. AEBSF, leupeptin, Lysogeny broth (LB) and Terrific Broth (TB) medium were purchased from Carl ROTH.

### 2.2. Cloning of Prdx2 and STAT3 DNA constructs

Prdx2 gene (codon-optimized for recombinant expression in *E. coli*) was cloned into the pD441 vector using the NEBuilder® HiFi DNA Assembly Master Mix (E2621, New England Biolabs, Ipswich, MA, USA). STAT3 gene for recombinant expression in *E. coli*, cloned into pET-SUMO, was kindly provided by the lab of Tobias Dick (German Cancer Research Center, Heidelberg, Germany). The core fragment STAT3 (CF-STAT3) gene was cloned into the pD441 vector using the NEBuilder® HiFi DNA Assembly Master Mix (E2621, New England Biolabs, Ipswich, MA, USA). All primers for the HiFi DNA Assembly were designed using Snapgene (version 5.2.2. GSL Biotech. San Diego, CA, USA) and are listed in **Supplementary Table 1**. Plasmids were purified using the PureYield™ Plasmid Miniprep System (A1222, Promega Biotech Ibérica S.L.).

### 2.3. Expression and purification of Prdx2 wild-type

His-SUMO-Prdx2 wild-type was expressed in *E. coli* NEB Turbo cells (C2984, New England Biolabs, Ipswich, MA, USA) in LB (50 μg/mL kanamycin). Cells were induced with 0.5 mM isopropyl β-D-1-thiogalactopyranoside (IPTG) (1758-1400, Inalco), grown overnight at 37°C, and harvested by centrifugation. Two protocols were used for Prdx2 purification: (1) where all buffers were supplemented with 2 mM DTT (reducing conditions); (2) where buffers were not supplemented with DTT (non-reducing conditions). Cells were lysed in 50 mM Tris-HCl pH 8.0, 500 mM NaCl, 0.1 mg/mL AEBSF, 0.1 μg/mL leupeptin, 50 μg/mL DNase I, 20 mM MgCl_2_ by sonication at 4°C, and the lysate was clarified at 39,000 *x g* at 4°C. The supernatant was incubated with pre-equilibrated Ni^2+^-NTA Sepharose High Performance beads (17526802, Cytiva, Marlborough, MA, USA) for 1 h at 4°C. Protein was eluted using a stepwise gradient of 50 mM Tris-HCl pH 8.0, 500 mM NaCl buffer and 500 mM imidazole. His-SUMO tags were cleaved by incubation with Ulp1 protease (1:40, w/w) during overnight dialysis in 50 mM Tris-HCl pH 8.0. The cleaved protein was recovered via reverse affinity chromatography with a final Superdex200 16/600 (28989335, Cytiva, Marlborough, MA, USA) step (50 mM Tris-HCl pH 7.6, 150 mM NaCl). Purified protein was stored at −80°C. Purity was assessed by SDS-PAGE and tandem liquid chromatography–mass spectrometry (LC-MS/MS). Buffers for reducing condition purification were Argon-flushed. The SEC buffer solution of non-reducing Prdx2 also contained 0.2 mM diethylenetriamine pentaacetate (DTPA).

### 2.4. Expression and purification of STAT3 wild-type and CF-STAT3 wild-type

Full-length His-SUMO-STAT3 wild-type and His-SUMO-core-fragment STAT3 wild-type (CF-STAT3, AA 136-689) were expressed in Rosetta(DE3)pLys or BL21(DE3) cells, respectively, in TB medium (50 μg/mL kanamycin and 34 µg/mL chloramphenicol for the Rosetta(DE3)pLys strain). Cells were induced with 1 mM IPTG (1758-1400, Inalco), grown overnight at 18°C, and harvested by centrifugation. Cells were lysed and purified by Ni-NTA affinity chromatography as in section 2.3, with all buffers containing 1 mM DTT. Full-length His-SUMO-STAT3 was further purified by HiScreen Capto ™ Q-Sepharose ion exchange (28926978, Cytiva, Marlborough, MA, USA). All proteins were loaded onto Superdex200 16/600 (28989335, Cytiva, Marlborough, MA, USA) (for CF) in 50 mM Tris-HCl pH 8, 150 mM NaCl, and 1 mM DTT, and stored at −80°C. Purity was assessed by SDS-PAGE and LC-MS/MS (only for full-length STAT3).

### 2.5. Annexin A2 wild-type production

His-Annexin A2 wild-type was expressed in HEK293F cells using the pcDNA3.1 vector. Secreted protein was purified via Ni^2+^-NTA Sepharose affinity (Cytiva, Marlborough, MA, USA) and eluted in 50 mM Tris-HCl pH 7.5, 500 mM NaCl buffer and 400 mM imidazole. Size-exclusion chromatography (SEC) (Superdex200 10/300 (Cytiva, Marlborough, MA, USA)) was performed as a last step in PBS, pH 7.4. Purity was assessed by SDS-PAGE and LC-MS/MS.

### 2.6. Peroxidase activity assay of Prdx2

The reaction was carried out in 96-well plates (249570, Polysorb, Nunc) at 25°C in 100 mM Hepes-NaOH pH 7.4, and 150 mM NaCl, 500 μM NADPH, 1 μM TrxR from *Corynebacterium glutamicum* (CgTrxR), and 8 μM Trx from *C. glutamicum* (CgTrx). Commercial recombinant human Prdx2 (Abcam, ab85331), or Prdx2 WT purified under reducing conditions were added at 0.5 µM final concentration, and the reaction was initiated by the addition of 20 µM H_2_O_2_. The decrease in absorbance at 340 nm was recorded every 30 s for 10 min using the Spectramax 340 PC (Molecular Devices). Negative control measurements were performed in the absence of H_2_O_2_. All data were normalized by subtracting corresponding negative controls as the background value and setting the time point when a decrease in absorbance started to be observed as time point zero.

### 2.7. Redox treatments

For complex reconstitution, proteins were pre-reduced with 50 mM DTT for 30 min at room temperature and desalted in the assay buffer. To oxidize Prdx2, Prdx2 was incubated with H_2_O_2_ (0.5-molar fold excess or 1.2-molar fold excess), and/or catalase (200 U/mL), or diamide (40-molar fold excess). Samples were incubated at room temperature.

### 2.8. Mass photometry

Samples were diluted (100 nM – 1 mM) in Hepes/NaOH or Tris/HCl and measured on Refeyn OneMP instrument (6,000 frames, 60 s, default settings). Calibration was performed using protein standards, i.e. β-amylase and IgG standards; or NativeMark™ Unstained Protein Standard (LC0725, ThermoFisher). Data were analyzed using default settings on DiscoverMP (version 2.1.1; Refeyn Ltd). Counts were binned and plotted vs. mass.

### 2.9. Size Exclusion Chromatography-Multi-Angle Light Scattering (SEC-MALS)

Pre-reduced Prdx2 and STAT3 were mixed at equimolar concentrations (1–50 μM) in 50 mM Tris-HCl pH 8, 15 mM NaCl in the absence and presence of H_2_O_2_ (10 µM), and analyzed using a Yarra SEC-3000 column (Phenomenex) connected to a HPLC Alliance system (Waters) equipped with a 2998 PDA detector (Waters), a TREOS II MALS detector (Wyatt Technology) and a RI-501 refractive index detector (Shodex). A bovine serum albumin sample (1 mg/mL) was used for data normalization. The Astra 7.3.0 software (Wyatt Technology) was used to analyze the data.

### 2.10. Analytical size exclusion chromatography (SEC)

Prdx2:STAT3 complex or individual proteins were loaded onto an analytical Superdex200 Increase 10/300 (GE28-9909-44) or 5/100 (GE28-9909-45) (Cytiva, Marlborough, MA, USA) in PBS, HEPES or Tris. Molecular weight standard proteins (Biorad) were also run under identical conditions. The molecular weights (MW) of the protein peaks were estimated by constructing a standard curve of log10(Mw) versus Kav, using the elution volumes and known molecular weights of the standards (K_av_ = (V_e_ - V_0_)/(V_t_ - V_0_), where V_e_ is the elution volume, V_0_ is the void volume, and V_t_ is the total volume. All values are in milliliters (mL) ^16^).

### 2.11. Negative-stain electron microscopy (EM)

Formvar/Carbon 400 Mesh, Cu grids (FCF400-Cu-50, Electron Microscopy Sciences) were glow discharged at 4-5 mA and 0.3 mbar vacuum for 30 s. 3 µL of freshly diluted sample (0.01-0.05 mg/mL) in the assay buffer was incubated on the grids for 30 s, followed by staining with 2% uranyl-acetate. Micrographs were collected on a JEOL 1400+ microscope, equipped with a LaB6 filament operating at 120 kV. Micrographs were recorded using a TVIPS F416 CCD camera using a nominal magnification of 60,000x, corresponding to a magnified pixel size of 1.94 Å/px and a defocus range of 0 to -1.5 µm.

### 2.12. Anaerobic conditions cryo-EM

The complex was reconstituted and isolated by SEC in an anaerobic atmosphere (<1 ppm O_2_, atmosphere of 3-4% hydrogen in nitrogen - Coy Laboratory Products). 3 µL of freshly isolated complex (0.3 mg/mL) was directly applied to freshly glow-discharged QUANTIFOIL R 1.2/1.3 Cu 300 (Q350CR1.3, Electron Microscopy Sciences) cryogenic electron microscopy (cryo-EM) grids. The grids were blotted using Whatman 595 blotting paper (Sigma-Aldrich) for 6 s at 4 °C and 100% humidity and then plunge-frozen into liquid ethane using a Vitrobot Mark IV (Thermo Fisher Scientific) under anaerobic conditions. A dataset was collected using a Titan Krios G4 equipped with a cold field-emission gun, Selectris X filter and Falcon 4 direct electron detector. EPU (version 3.2) was used for automated data acquisition of 12,199 movies in electron event representation (EER) format. Movies were collected with a pixel size of 0.573 Å and a total dose of 60 *e*^−^/Å^2^ contained in 1,098 EER frames. The defocus ranged from −0.6 to −2.2 μm.

### 2.13. Electron microscopy (EM) data processing

For negative staining EM data processing, the micrographs were processed using CryoSparc v4.6.0 ^17^. After running patch CTF estimation, particles were picked by blob-picker and extracted using a box size of 256 px. These particles were subjected to 2D classification, resulting in 1,583 final particles. Cryo-EM data was analyzed in CryoSPARC (v4.6.0) ^17^. After the micrographs were imported, they were motion corrected with Patch Motion Correction. The CTF was estimated with Patch CTF estimation. Particles were picked with Topaz (cite) and extracted in a 1,260 pixel box, downscaled to 216 pixels. The particle stack was cleaned up with multiple rounds of 2D classification resulting in 592,392 final particles.

### 2.14. Crosslinking and complex stabilization

Crosslinking was performed using glutaraldehyde (0.03–1%) or CovalX (W2010k100 and W2010k200, CovalX) (0.01–0.1 mg/mL) reagents for 1 or 2 h at room temperature, respectively. Oxidized Prdx2 (0.5-fold excess H_2_O_2_, followed by 200 U/mL catalase incubation) and freshly pre-reduced CF-STAT3 WT were mixed in a 2:1 molar ratio (8 mM:4 mM). Reactions were quenched with 50 mM Tris/HCl pH 8.

### 2.15. Mass spectrometry tandem Liquid chromatography–mass spectrometry (LC-MS/MS)

Peptide separation was performed using a reversed-phase analytical column (Acclaim PepMap RSLC, 0.075 × 250 mm, Thermo Scientific) (linear gradient 4%–27.5% solvent B (0.1% formic acid (FA) in 98% acetonitrile (ACN)) for 60 min, 27.5%–40% solvent B for 10 min, 40%–95% solvent B for 1 min and holding at 95% for 10 min, 300 nL/min) on an Ultimate 3000 RSLC system. Peptides were analyzed by an Orbitrap Fusion Lumos tribrid mass spectrometer (ThermoFisher Scientific) and subjected to NSI source followed by tandem mass spectrometry (MS/MS) in Fusion Lumos coupled online to the nano-LC. Intact peptides were detected in the Orbitrap at a resolution of 120,000 and selected for MS/MS using HCD setting at 30; ion fragments were detected in the Orbitrap at a resolution of 30,000. A data-dependent procedure (one MS scan followed by MS/MS scans) was applied for 3 s for ions above a threshold ion count of 2.0E4 in the MS survey scan with 40.0 s dynamic exclusion (electrospray voltage of 2.1 kV). MS1 spectra were obtained with an AGC target of 4E5 ions and a maximum injection time of 50 ms, MS2 spectra were acquired with an AGC target of 5E4 ions and a maximum injection time set to dynamic (m/z scan range 375–1800). MS/MS data was processed using Sequest HT search engine within Proteome Discoverer 2.5 SP.

### 2.16. Matrix-Assisted Laser Desorption/Ionization Time-of-Flight Mass Spectrometry (MALDI-TOF MS)

Protein samples were exchanged into 0.1% *v/v* trifluoroacetic acid (TFA); alkylated samples were obtained by treatment with 12.5 mM iodoacetamide (IAM) for 20 min in the dark and re-buffering to 0.1% *v/v* TFA. 1:1:1 mixture containing protein, 2,5-dihydroxyacetophenone (DHAP) matrix (8231829, Bruker), and 2% *v/v* TFA was spotted in duplicate onto an MTP ground steel plate (8280784, Bruker). Spectra were acquired on a Ultraflextreme enhanced MALDI TOF/TOF-MS system (Bruker) in linear positive mode (range of 5 000 to 50 000 m/z), processed in FlexAnalysis; duplicates per sample. All acquisition methods were provided by the manufacturer and optimized and calibrated with in-house made calibration standard (22 to 44 kDa, 4 calibrants).

## 3. Results

### 3.1. Prdx2 purifies as decamers, while full-length STAT3 and annexin A2 yield stable monomers

To reconstitute the Prdx2:STAT3 redox-relay complex *in vitro*, proteins were expressed and purified. We initially attempted to purify untagged Prdx2, but it resulted in contamination with *E. coli* AhpC (data not shown). Hence, we resorted to purifying Prdx2 with an N-terminal His-SUMO-tag to enhance solubility and yield, and purified under reducing conditions with DTT ^18^ (**Fig. S1A**). After His-SUMO cleavage, Prdx2 retained its native decameric assembly in reducing conditions, as confirmed by size-exclusion chromatography (SEC), mass photometry, and negative-staining electron microscopy (EM) (**Fig. 2A-C)**. Activity was validated by an NADPH-coupled peroxidase assay using H_2_O_2_, showing peroxidase function (**Fig. 2D and Fig. S1D)**. Notably, the presence of the N-terminal SUMO-tag disrupted the decamer formation (**Fig. 2B-C**), highlighting the importance of tag removal for structural integrity. Although Ulp1 digestion is generally performed under reducing conditions, we achieved efficient cleavage under non-reducing conditions (**Fig. S1B**).

**Figure 2.**
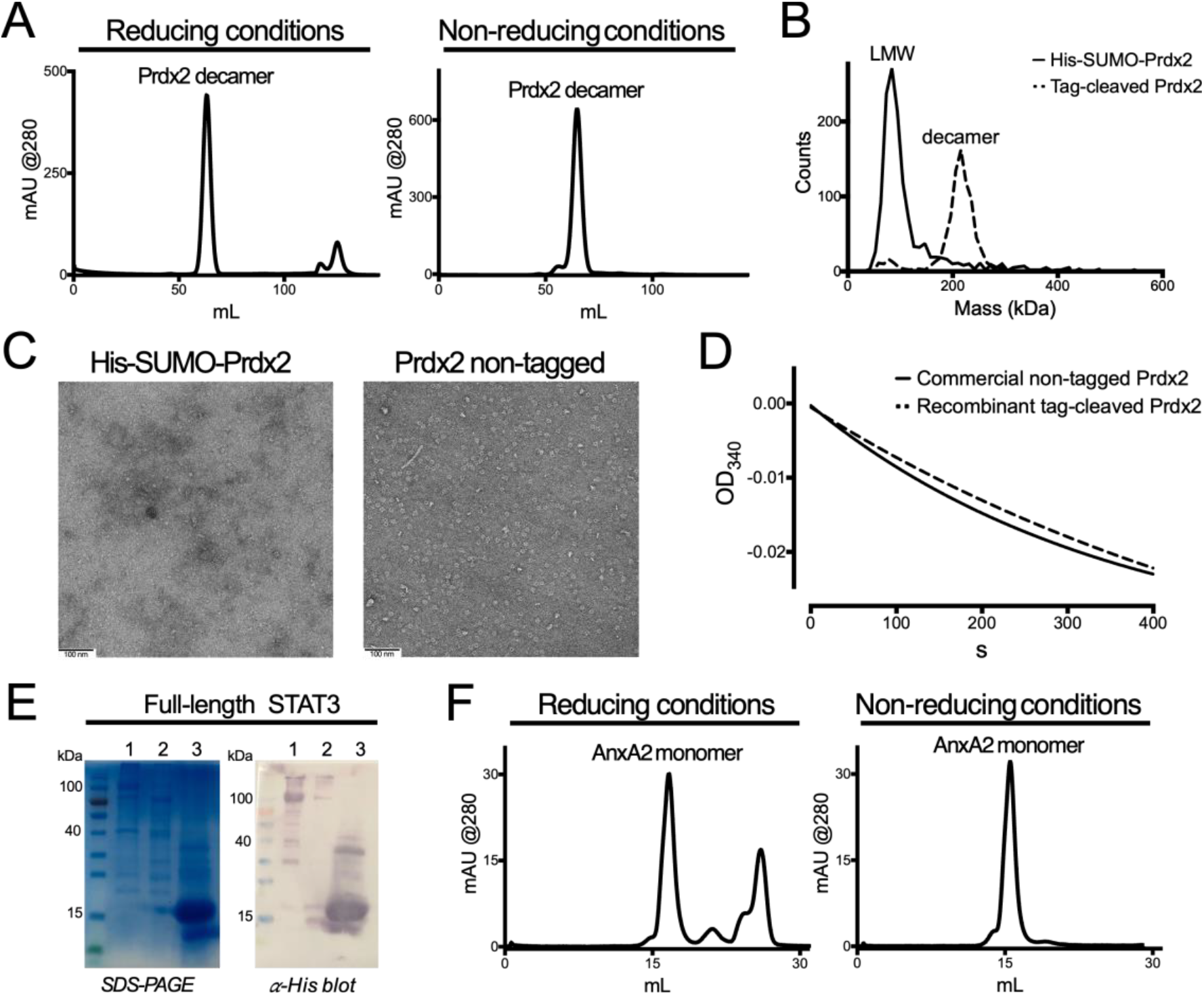
Prdx2 purifies as active decamers, while full-length STAT3 and AnxA2 yield stable monomers. **(A)** SEC profiles of Prdx2 purified in reducing (presence of DTT) and non-reducing (absence of DTT) conditions show a predominant decameric peak consistent with known oligomeric state. **(B)** Mass photometry data shows that His-SUMO-Prdx2 primarily exists as low molecular weight (LMW) oligomers, whereas tag-cleaved Prdx2 forms decamers under reducing conditions. Measurements were conducted using diluted stock solutions at 100 nM. **(C)** Negative-stain electron microscopy (EM) reveals that Prdx2 remains in a LMW oligomeric state when fused to the His-SUMO-tag. When the His-SUMO tag is cleaved, Prdx2 forms decamers. Micrographs were acquired under reducing conditions at a protein concentration of 0.01 mg/mL. Full-picture on Fig. S2. **(D)** Time-course of NADPH oxidation by Prdx2 purified under reducing conditions shows that Prdx2 retains peroxidase activity following His-SUMO cleavage. **(E)** 12% SDS-PAGE and western blot (anti-His) analysis of different populations (1, 2, 3) from SEC show degradation products of full-length STAT3 (cropped gel/blot image). **(F)** SEC profiles of purified Annexin A2 in reducing and non-reducing conditions show that AnxA2 is mainly monomeric.

By contrast, recombinant full-length STAT3 expressed in bacteria showed poor stability. Despite His-SUMO tagging and purification by affinity chromatography, ion exchange, and SEC, degradation was observed after SEC (**Fig. 2E**). A small intact STAT3 with minor C-terminal degradations was nevertheless retained for downstream studies. The final component, AnnexinA2 (AnxA2), was expressed with an N-terminal His-TEV in HEK293F and purified via affinity chromatography and SEC. AnxA2 was obtained in pure, mainly monomeric form regardless of redox state (**Fig. 2F and Fig. S1C**).

### 3.2. Prdx2 and STAT3 fail to form a stable complex

To assess whether Prdx2 and full-length STAT3 form a stable complex *in vitro*, as reported in cells ^10,11^, we mixed the two proteins at a 1:1 ratio and performed SEC-MALS ^19^ and mass photometry ^20^ under low salt conditions. Both methods showed Prdx2 and STAT3 eluting as distinct species, with no complex detected even after oxidative treatment with H_2_O_2_ (SEC-MALS data not shown, and **Fig. 3**). Because AnxA2 has been proposed to scaffold the Prdx2:STAT3 interaction in cells ^11^, we repeated the assays in its presence. However, no defined Prdx2:STAT3 complex was formed, and EM revealed aggregation rather than specific assemblies (**Fig. S2**). Overall, biochemical assays do not support stable Prdx2–STAT3 complex formation *in vitro*, and AnxA2 does not appear sufficient to stabilize the interaction.

**Figure 3.**
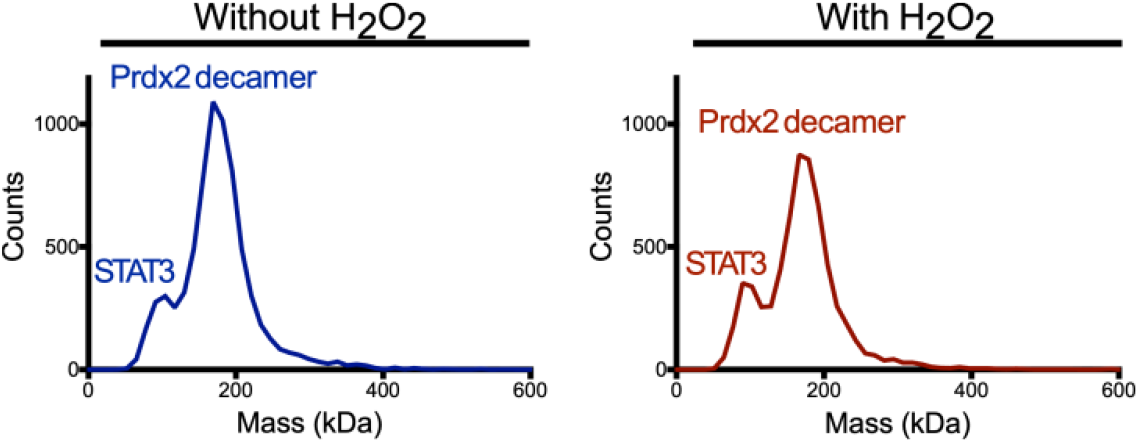
Prdx2:STAT3 complex is detected neither in the absence nor presence of H_2_O_2_ or AnxA2. **(A)** Mass photometry analysis of Prdx2 and full-length STAT3 in complex in the absence and presence of H_2_O_2_ reveal discrete, non-overlapping elution profiles consistent with decameric Prdx2 and monomeric STAT3. Pre-reduced proteins were mixed in a 1:1 molar ratio (1 mM total protein) in low salt buffer with or without 10-molar excess of H_2_O_2_. Two peaks were consistently observed: 101 kDa and 175 kDa without H_2_O_2_ and 99 kDa and 174 kDa with H_2_O_2_, corresponding to full-length His-SUMO-STAT3 and Prdx2 decamer, respectively. No additional peaks indicative of complex formation were detected.

### 3.3. Formation of stable disulfide-linked Prdx2:STAT3 complexes using CF-STAT3 under anaerobic conditions

To test whether improving the quality of STAT3 would increase the chances of complex formation, we turned to a core-fragment variant of STAT3 (CF-STAT3, AA 136–689), which lacks disordered termini but retains functional domains for dimerization and DNA binding ^21-24^ (**Fig. S3A**). This construct, previously crystallized (PDB IDs: 6TLC, 6QHD, 6NJS), expressed efficiently as a His-SUMO–tagged protein, showed no degradation, and eluted as a stable monomer (∼76 kDa; **Fig. S3B-C)**. However, co-incubation in low salt buffer with Prdx2 in a 1:1 ratio with Tween20 did not result in complex formation when assessed by analytical SEC (**Fig. 4A**). Tween-20 was included to mimic membrane-associated environments where the interaction has been reported ^11^.

**Figure 4.**
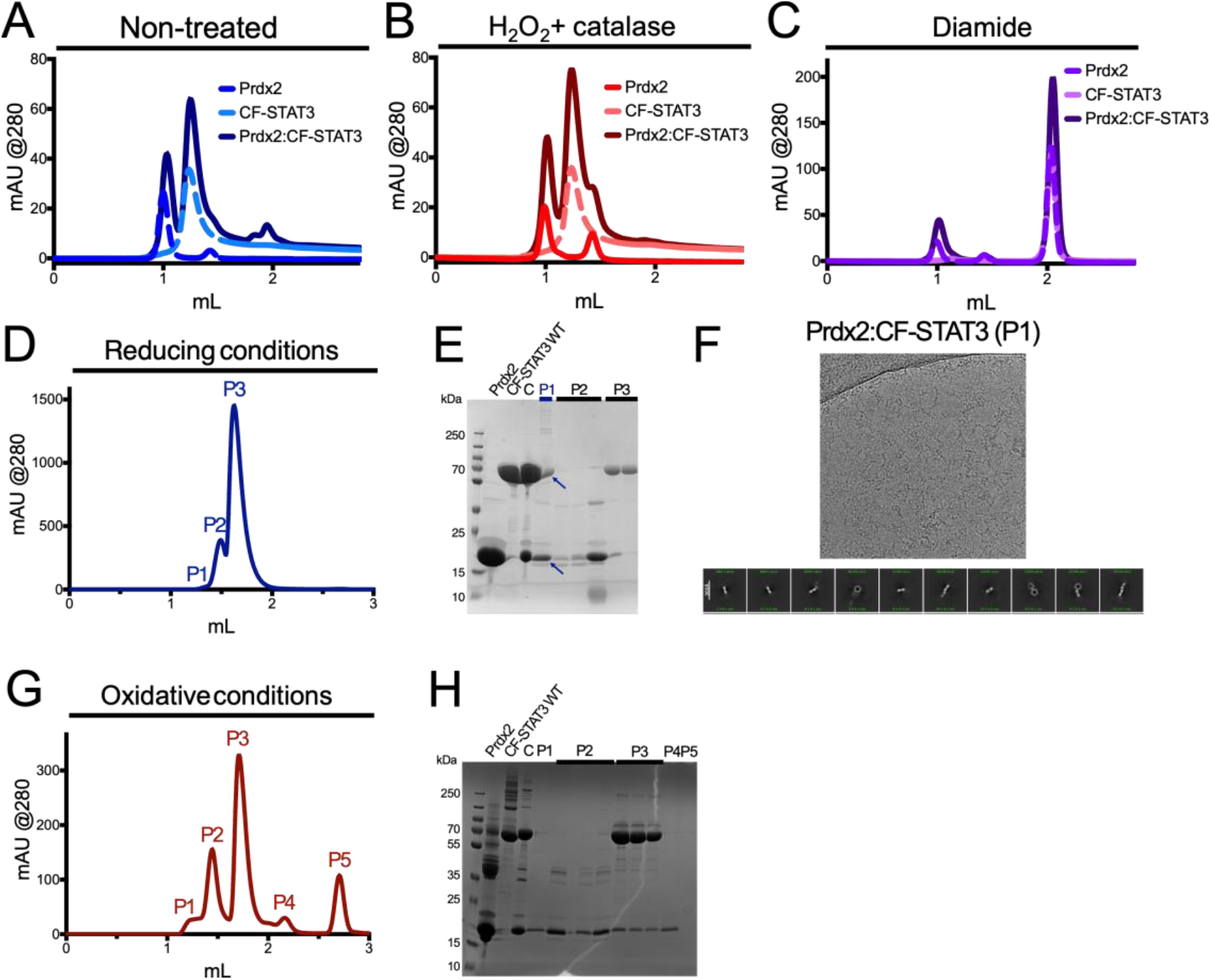
Absence of a mixed Prdx2:CF-STAT3 population indicates lack of complex formation. **(A, B and C)** SEC profile of Prdx2 reveals a population consistent with decamers (∼220 kDa), whereas of CF-STAT3 is detected as a monomer (∼90 kDa). Mixture of Prdx2 and CF-STAT3 under oxidizing conditions fails to yield intermediate or shifted mass species, supporting the absence of stable Prdx2:CF-STAT3 complex formation in vitro. Pre-reduced proteins were mixed in a 1:1 molar ratio in 25 mM Tris-HCl pH 7.4, 50 mM NaCl, and 0.05% Tween20, with 0.5-fold molar excess of H_2_O_2_ plus 200 u/mL catalase, or with 40-fold molar excess of diamide. **(D)** Chromatogram of SEC under anaerobic conditions and **(E)** 12% SDS-PAGE gel (non-reducing conditions) show Prdx2:CF-STAT3 complex isolation in the presence of reducing agent under anaerobic conditions. Prdx2 and CF-STAT3 were mixed in a 1:1 molar ratio (70 mM each protein) in 50 mM Tris-HCl pH 7.4, and 150 mM NaCl with 5 mM DTT under controlled anaerobic conditions. Peak 1 presents a mixture of Prdx2 and CF-STAT3 (blue arrows) (cropped gel image). C: reconstituted complex; P: peaks from the SEC. **(F)** Cryo-EM micrograph and 2D class averages of selected particles show that only Prdx2 particles are present as isolated particles, indicating that Prdx2:CF-STAT3 isolated complex is not stable (full-picture on Fig. S4). **(G)** SEC chromatogram obtained under anaerobic conditions and **(H)** 12% SDS-PAGE gel (non-reducing conditions) show no stable disulfide-linked Prdx2:CF-STAT3 complex under oxidative, anaerobic conditions. Prdx2 and CF-STAT3 were mixed in a 2:1 molar ratio in 50 mM Tris-HCl pH 7.4, and 150 mM NaCl with 0.5-molar excess H_2_O_2_ and 200 U/mL catalase, and 100-molar excess of maleimide under controlled anaerobic conditions (cropped gel image). C: reconstituted complex; P: peak from the SEC.

We next tested oxidative treatments, reasoning that disulfide exchange might promote stable complex formation as seen in cells ^11,25-27^. Prdx2 and CF-STAT3 were exposed to H_2_O_2_ (0.5-fold excess) followed by catalase or to diamide (40-fold excess) in the presence of Tween-20 ^28^ (**Fig. 4B-C**) ^25-27^. However, SEC consistently showed only separate species, indicating no stable assembly under these conditions. These findings suggested that the transient nature of the Prdx2:CF-STAT3 interaction and the sensitivity of cysteines to overoxidation may hinder reconstitution *in vitro*.

To overcome these issues, we tested anaerobic conditions (<1 ppm O_2_, atmosphere of 3-4% H_2_ in N_2_). Prdx2 and CF-STAT3 were incubated under reducing (5 mM DTT) or oxidative conditions (0.5-fold excess H_2_O_2_, followed by 200 U/mL catalase incubation and 100-molar fold excess diamide). Analytical SEC and SDS-PAGE under reducing conditions suggested possible complex formation (**Fig. 4D**), prompting cryo-EM analysis with grids prepared under anaerobic conditions. However, non-reducing SDS-PAGE showed no disulfide-linked species (**Fig. 4E**) and cryo-EM imaging revealed only Prdx2 decamers without STAT3 density (**Fig. 4F**). Attempts to stabilize interactions by oxidative trapping and thiol alkylation also failed (**Fig. 4G-H**).

### 3.4. Chemical crosslinkers show potential in complex stabilization

Finally, we tested chemical crosslinking to stabilize the Prdx2:STAT3 complex. Two crosslinkers were tested: glutaraldehyde and the CovalX high-mass MALDI crosslinker. Titration experiments revealed that CovalX efficiently generated high–molecular weight species containing both Prdx2 and CF-STAT3, consistent with complex formation, whereas glutaraldehyde caused mainly nonspecific aggregation and was not pursued further (data not shown). A concentration of 0.1 mg/mL CovalX was selected as it minimized free protein while preserving complex integrity. Crosslinked species were isolated by Superdex200 Increase 10/300 and analyzed by SDS-PAGE and western blotting **(Figure 5A-B**). Negative-stain EM confirmed the homogeneity and stability of the crosslinked complex (**Figure 5C**). Building on this, we optimized cryo-EM grid preparation—adjusting graphene oxide grid quality, plunge-freezing parameters, and protein concentration—using a JEOL JEM-1400 microscope. However, crosslinking proved inconsistent in later experiments, underscoring the inherent instability of the Prdx2:STAT3 complex.

**Figure 5.**
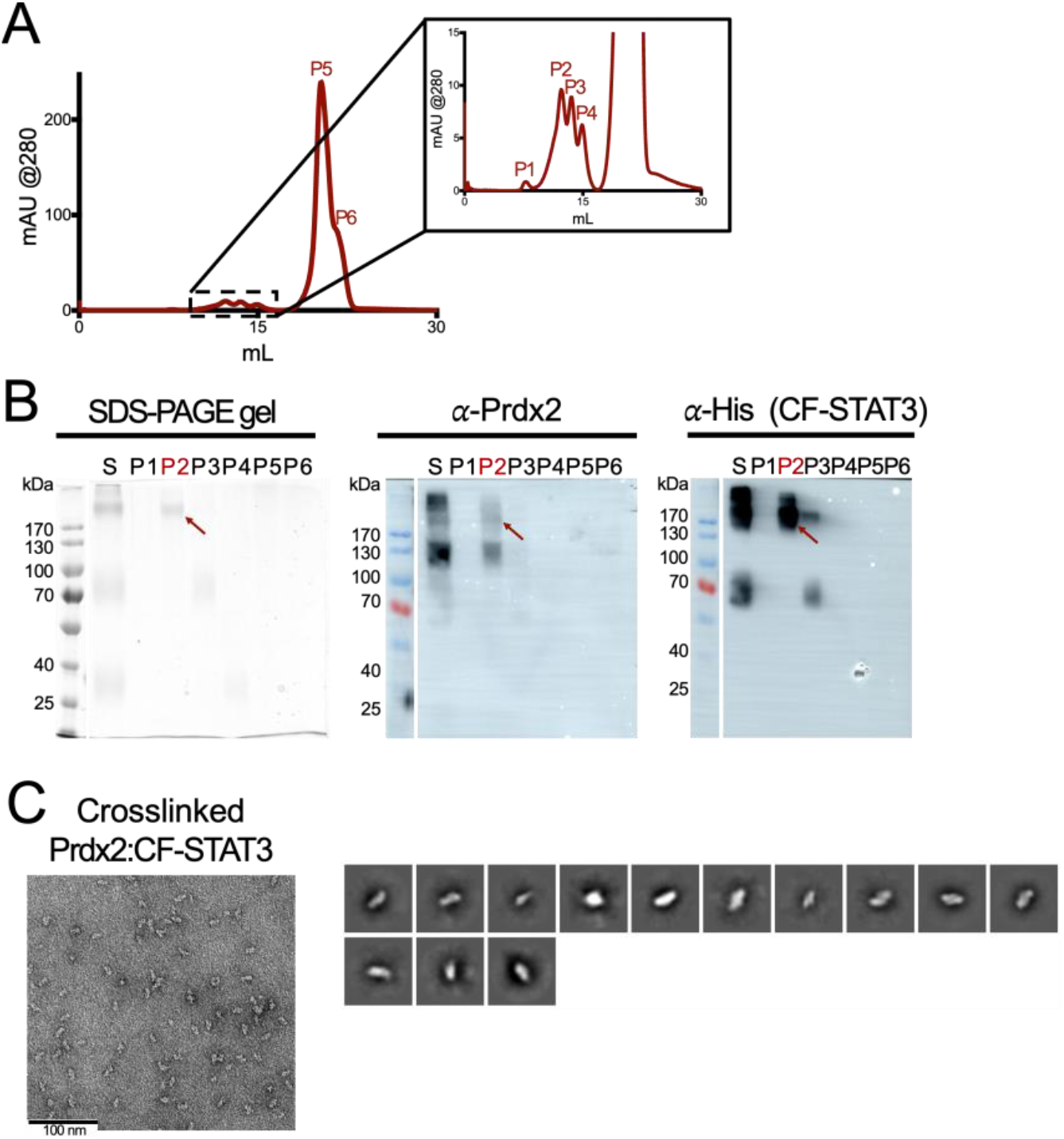
Oxidized Prdx2 and reduced CF-STAT3 form a stable complex when treated with chemical crosslinkers. **(A)** Chemical crosslinkers, i.e. CovalX. CovalX K100 at 0.1 mg/mL effectively stabilizes the Prdx2:CF-STAT3 complex, producing a distinct peak corresponding to ∼440 kDa (sample P2). **(B)** 12% SDS-PAGE of the respective elution fractions of panel A in non-reducing conditions, immunoblot with anti-Prdx2 antibody and immunoblot with anti-His antibody detecting His-SUMO-CF-STAT3, recognize Prdx2 and CF-STAT3 in the peak 2 sample from the SEC (red arrows) (cropped gel/blot image). S: sample of reconstituted complex; P: peaks from the SEC. **(C)** Representative negative-stain electron microscopy (EM) image and 2D class averages from negative stain EM data show that crosslinked Prdx2:CF-STAT3 forms a homogeneous and stable population under oxidative conditions. Micrographs were acquired under reducing conditions at a protein concentration of 0.04 mg/mL.

### 3.5. Prdx2 hyperoxidation under reducing conditions prevents complex formation with CF-STAT3

Building on our observation that complex formation was only detectable under anerobic conditions, we hypothesized that Prdx2 hyperoxidation blocks its interaction with CF-STAT3. Indeed, Prdx2 proved highly sensitive to hyperoxidation at its peroxidatic cysteine (Cys51), a residue essential for both catalysis and redox-relay activity.

Prdx2, purified and stored with DTT gradually lost activity, and MALDI-TOF MS showed conversion of Cys51 to sulfinic or sulfonic acids forms, modifications that abolish catalytic function and prevent disulfide bond formation (**Figure 6A**). Even after further DTT treatment, the oxidized mass shift persisted, confirming irreversible hyperoxidation (data not shown). Excess H_2_O_2_ exposure produced a similar effect (**Table 1**), indicating that both reductive and oxidative stress can push Prdx2 beyond its functional redox window, thereby preventing transient complex formation with CF-STAT3.

**Figure 6.**
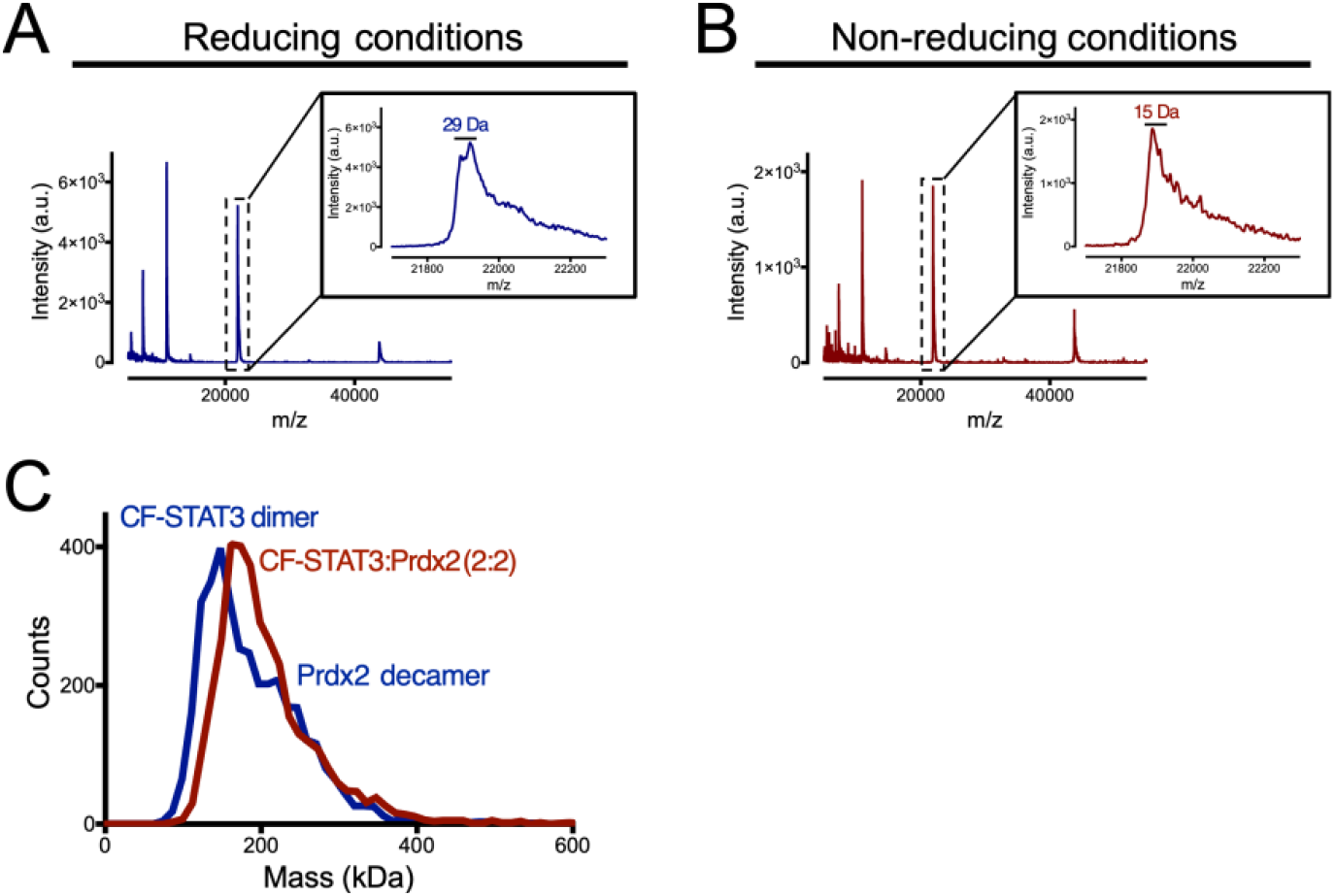
Prdx2 purified and stored without any reducing agent does not form the hyperoxidized state. **(A and B)** MALDI-TOF MS data reveal that Prdx2 purified and stored under reducing conditions presents hyperoxidation in the sulfinic acid form, whereas Prdx2 purified in the absence of any reducing agents does not present hyperoxidation. When stored in the presence of 2 mM DTT (reducing conditions), Prdx2 displays a population corresponding to the sulfinic acid form, indicated by a mass shift of +32 Da (two extra oxygens). In contrast, Prdx2 purified and stored under non-reducing conditions shows a population with a +16 Da shift (one extra oxygen), consistent with the sulfenic acid form, indicating no hyperoxidation. **(C)** Oxidized Prdx2 forms transient complexes with CF-STAT3. Mass photometry of oxidized Prdx2 mixed with CF-STAT3 at 1 μM under non-reducing conditions shows a small fraction of intermediate mass species (∼190 kDa), suggesting a weakly associated Prdx2:STAT3 population consistent with a 2:2 ratio. Additional peaks corresponding to dimeric CF-STAT3 and decameric Prdx2 were also detected under reducing conditions. Measurements were conducted using diluted stock solutions at 1 μM.

**Table 1.** The peroxidatic Cys of Prdx2 (Cys51) is primarily overoxidized to the sulfinic acid (Cys_P_-SO_2_H) form after treatment with a 1.2-molar excess of H_2_O_2_. Prdx2 was incubated with 1.2-molar excess of H_2_O_2_, 250-molar excess of dimedone and 100-molar excess of iodoacetamide (IAM). Proteins were analyzed by LC-MS/MS.

|  | -SH | -SOH | -SO<sub>2</sub>H | -SO<sub>3</sub>H |
| --- | --- | --- | --- | --- |
| **Cys51** | 6.8% | 3.2% | 84.5% | 4.5% |
| **Cys70** | 89.9% | 0.6% | 7.9% | 1.5% |
| **Cys176** | 91.4% | 4.4% | 2.6% | 1.7% |

Consistent with published protocols ^29-32^, we found that omitting DTT and including the metal chelator DTPA during purification (“non-reducing conditions”) preserved Cys51 in its functional sulfenic or disulfide state (**Figure 6B**). Under these conditions, Prdx2 remained catalytically competent and capable of disulfide exchange.

As hypothesized, Prdx2 prepared in this way supported a reproducible, redox-sensitive interaction with CF-STAT3. Mass photometry revealed a mixed-population corresponding to Prdx2:CF-STAT3 complexes (**Figure 6C**). Although transient and low in abundance, this interaction was consistently observed, underscoring the importance of maintaining Prdx2 in its proper redox state for redox-relay engagement.

These results highlight the critical role of redox control during protein purification and handling, and they provide an experimental framework for further structural and biochemical analysis of Prdx2-mediated signaling, including strategies to stabilize or capture transient relay complexes.

## 4. Discussion

This study highlights the challenges of reconstituting transient redox-relay complexes such as Prdx2:STAT3 *in vitro*. Unlike stable protein-protein interactions, redox relays rely on short-lived thiol-disulfide exchange intermediates that are highly sensitive to the redox environment and protein conformation ^7-11^. We were unable to capture a stable Prdx2:STAT3 complex under standard biochemical conditions, underscoring the need to maintain the native redox state of cysteines in both proteins.

A key observation is that DTT promotes Prdx2 hyperoxidation, which inactivates its peroxidatic cysteine (Cys51) and prevents disulfide exchange with STAT3 ^1-4^. DTT may continuously recycle Prdx2 to its reduced form, leaving the peroxidatic cysteine repeatedly exposed to H_2_O_2_ and thereby accelerating hyperoxidation to sulfinic or sulfonic acid forms, rendering Prdx2 catalytically inactive (**Fig. S1A, Fig. 6A and Fig. 3**). Prdx2 purified under non-reducing conditions stabilized Cys51 in its functional state (**Fig. S1B and Fig. 6B**), enabling transiently interaction with CF-STAT3 (**Fig. 6C**). This suggests that prior failures to detect Prdx2:STAT3 relays may have at least partially stemmed from unrecognized hyperoxidation.

CF-STAT3 proved more stable than full-length STAT3 (**Fig. S3**), though stable complex formation was not achieved (**Fig. 4A-C**). Future work could employ CF-STAT3 cysteine mutants to favor redox-relay interactions over alternative STAT3 oligomerization pathways ^33,34^. Also, addition of AnxA2 did not stabilize the complex ^11^, likely reflecting the absence of additional cellular factors (e.g., metals, modifications) required for its scaffolding role *in vivo*. EM revealed heterogeneity and aggregation (**Fig. S2**), consistent with context-dependent, transient interactions.

Two stabilizing strategies were tested. Under anaerobic conditions, reducing environments suggested complex formation, but oxidative treatments did not yield stable disulfide-linked species (**Fig. 4D-H**). Chemical crosslinking with CovalX was more effective, producing a homogeneous, crosslinked complex suitable for EM (**Fig. 5**), though reproducibility remained limited.

In summary, Prdx2:STAT3 assembly *in vitro* is hindered by weak, transient interactions and strict redox dependence. These findings are consistent with Langford *et al*. (2018), who predicted that Prdx2– STAT3 reactions are kinetically disfavored in the cytosol ^35^, and with Honzejkova *et al*. (2024), who reported similar difficulties stabilizing ASK1:Trx1 for cryo-EM ^36^. Our results emphasize that buffer composition, redox control, and protein quality are critical for redox-relay studies. Future approaches such as site-specific crosslinkers, rapid-mixing kinetics, or redox-trapped cryo-EM may provide the structural insights needed to fully understand Prdx2-mediated signaling.

## Supplementary Materials

The following supporting information can be downloaded at: https://www.mdpi.com/article/doi/s1, Table S1. List of primers used in this study; Table S2. Intersubunit hydrogen bonds from the AlphaFold 3 predicted complex between dimeric Prdx2 and STAT3; Table S3. Intersubunit hydrophobic interactions from the AlphaFold 3 predicted complex between dimeric Prdx2 and STAT3; Table S4. Intersubunit salt bridges from the AlphaFold 3 predicted complex between dimeric Prdx2 and STAT3; Table S5. Intersubunit hydrogen bonds from the AlphaFold 3 predicted complex between dimeric Prdx2, AnxA2, and STAT3; Table S6. Intersubunit hydrophobic interactions from the AlphaFold 3 predicted complex between dimeric Prdx2, AnxA2, and STAT3; Table S7. Intersubunit salt bridges from the AlphaFold 3 predicted complex between dimeric Prdx2, AnxA2, and STAT3; Figure S1. Proteins are successfully purified to homogeneity; Figure S2. Negative-stain EM images of Prdx2:STAT3 mixtures reveal diffuse, heterogeneous particles with no evidence of higher-order assemblies; Figure S3. CF-STAT3 does not show degradation products; Figure S4. 2D class averages of selected particles show that only Prdx2 particles are present as isolated particles.

## Author Contributions

Conceptualization – J.M.P., E.B., D.E., and J.M.; Data curation – J.M.; Formal analysis – J.M.P., E.B., K.W., V.R.A.D., S.J., and D.V.; Funding acquisition – J.M.P., B.J.M., and J.M.; Investigation – J.M.P., E.B., K.W., V.R.A.D., S.J., and D.V.; Methodology – J.M.P., E.B., K.W., V.R.A.D., S.J., and D.V.; Project administration – J.M.P. and J.M.; Resources – B.J.M., and J.M.; Supervision – E.B., B.J.M., D.E., and J.M.; Visualization – J.M.P., and J.M.; Writing – original draft – J.M.P., E.B., D.E., and J.M.; Writing – review and editing – J.M.P., E.B., K.W., V.R.A.D., S.J., D.V., B.J.M., D.E., and J.M.. All authors have read and agreed to the published version of the manuscript.

## Institutional Review Board Statement

Not applicable.

## Informed Consent Statement

Not applicable.

## Data Availability Statement

The data supporting the findings of this study are available from the corresponding authors upon reasonable request.

## Acknowledgments

We thank Dr. Christoph W. Müller for insightful guidance on STAT3 design. We also acknowledge the VIB Protein Core Facility and the EMBO Practical Course on Integrative Structural Biology for their technical support. We thank the VIB-VUB facility for Biological Electron Cryogenic Microscopy (BECM), especially Dr. Marcus Fislage and Dr. Dirk Reiter, for the support in sample preparation and data analysis, and gratefully acknowledge the Central Electron Microscopy Facility at the Max Planck Institute of Biophysics for providing access to cryo-EM instrumentation and expert technical support. We would like to thank Dr. Thomas Zögg for his expertise in the use of chemical crosslinkers and providing the CovalX kit. The authors have reviewed and edited the output and take full responsibility for the content of this publication.

## References

1 Perkins, A., Poole, L. B. & Karplus, P. A. Tuning of peroxiredoxin catalysis for various physiological roles. Biochemistry 53, 7693–7705, doi:10.1021/bi5013222 (2014).

2 Nelson, K. J. et al. Analysis of the peroxiredoxin family: using active-site structure and sequence information for global classification and residue analysis. Proteins 79, 947–964, doi:10.1002/prot.22936 (2011).

3 Rhee, S. G. & Kil, I. S. Multiple Functions and Regulation of Mammalian Peroxiredoxins. Annu Rev Biochem 86, 749–775, doi:10.1146/annurev-biochem-060815-014431 (2017).

4 Wood, Z. A., Schroder, E., Robin Harris, J. & Poole, L. B. Structure, mechanism and regulation of peroxiredoxins. Trends Biochem Sci 28, 32–40, doi:10.1016/s0968-0004(02)00003-8 (2003).

5 Baker, L. M., Raudonikiene, A., Hoffman, P. S. & Poole, L. B. Essential thioredoxin-dependent peroxiredoxin system from Helicobacter pylori: genetic and kinetic characterization. J Bacteriol 183, 1961–1973, doi:10.1128/JB.183.6.1961-1973.2001 (2001).

6 Poole, L. B. The catalytic mechanism of peroxiredoxins. Subcell Biochem 44, 61–81, doi:10.1007/978-1-4020-6051-9_4 (2007).

7 Stocker, S., Maurer, M., Ruppert, T. & Dick, T. P. A role for 2-Cys peroxiredoxins in facilitating cytosolic protein thiol oxidation. Nat Chem Biol 14, 148–155, doi:10.1038/nchembio.2536 (2018).

8 Jarvis, R. M., Hughes, S. M. & Ledgerwood, E. C. Peroxiredoxin 1 functions as a signal peroxidase to receive, transduce, and transmit peroxide signals in mammalian cells. Free Radic Biol Med 53, 1522–1530, doi:10.1016/j.freeradbiomed.2012.08.001 (2012).

9 Vo, T. N. et al. Prdx1 Interacts with ASK1 upon Exposure to H2O2 and Independently of a Scaffolding Protein. Antioxidants (Basel) 10, doi:10.3390/antiox10071060 (2021).

10 Sobotta, M. C. et al. Peroxiredoxin-2 and STAT3 form a redox relay for H2O2 signaling. Nat Chem Biol 11, 64–70, doi:10.1038/nchembio.1695 (2015).

11 Talwar, D., Messens, J. & Dick, T. P. A role for annexin A2 in scaffolding the peroxiredoxin 2-STAT3 redox relay complex. Nat Commun 11, 4512, doi:10.1038/s41467-020-18324-9 (2020).

12 Hirano, T., Ishihara, K. & Hibi, M. Roles of STAT3 in mediating the cell growth, differentiation and survival signals relayed through the IL-6 family of cytokine receptors. Oncogene 19, 2548–2556, doi:10.1038/sj.onc.1203551 (2000).

13 Laudisi, F., Cherubini, F., Monteleone, G. & Stolfi, C. STAT3 Interactors as Potential Therapeutic Targets for Cancer Treatment. Int J Mol Sci 19, doi:10.3390/ijms19061787 (2018).

14 Laudisi, F. et al. Progranulin sustains STAT3 hyper-activation and oncogenic function in colorectal cancer cells. Mol Oncol 13, 2142–2159, doi:10.1002/1878-0261.12552 (2019).

15 Goemaere, J. & Knoops, B. Peroxiredoxin distribution in the mouse brain with emphasis on neuronal populations affected in neurodegenerative disorders. J Comp Neurol 520, 258–280, doi:10.1002/cne.22689 (2012).

16 O’Fagain, C., Cummins, P. M. & O’Connor, B. F. Gel-Filtration Chromatography. Methods Mol Biol 1485, 15–25, doi:10.1007/978-1-4939-6412-3_2 (2017).

17 Punjani, A., Rubinstein, J. L., Fleet, D. J. & Brubaker, M. A. cryoSPARC: algorithms for rapid unsupervised cryo-EM structure determination. Nat Methods 14, 290–296, doi:10.1038/nmeth.4169 (2017).

18 Engelman, R. et al. Multilevel regulation of 2-Cys peroxiredoxin reaction cycle by S-nitrosylation. J Biol Chem 288, 11312–11324, doi:10.1074/jbc.M112.433755 (2013).

19 Some, D., Amartely, H., Tsadok, A. & Lebendiker, M. Characterization of Proteins by Size-Exclusion Chromatography Coupled to Multi-Angle Light Scattering (SEC-MALS). J Vis Exp, doi:10.3791/59615 (2019).

20 Soltermann, F. et al. Quantifying Protein-Protein Interactions by Molecular Counting with Mass Photometry. Angew Chem Int Ed Engl 59, 10774–10779, doi:10.1002/anie.202001578 (2020).

21 Becker, S., Corthals, G. L., Aebersold, R., Groner, B. & Muller, C. W. Expression of a tyrosine phosphorylated, DNA binding Stat3beta dimer in bacteria. FEBS Lett 441, 141–147, doi:10.1016/s0014-5793(98)01543-9 (1998).

22 La Sala, G. et al. Selective inhibition of STAT3 signaling using monobodies targeting the coiled-coil and N-terminal domains. Nat Commun 11, 4115, doi:10.1038/s41467-020-17920-z (2020).

23 Belo, Y. et al. Unexpected implications of STAT3 acetylation revealed by genetic encoding of acetyl-lysine. Biochim Biophys Acta Gen Subj 1863, 1343–1350, doi:10.1016/j.bbagen.2019.05.019 (2019).

24 Bai, L. et al. A Potent and Selective Small-Molecule Degrader of STAT3 Achieves Complete Tumor Regression In Vivo. Cancer Cell 36, 498–511 e417, doi:10.1016/j.ccell.2019.10.002 (2019).

25 Biswas, S., Chida, A. S. & Rahman, I. Redox modifications of protein-thiols: emerging roles in cell signaling. Biochem Pharmacol 71, 551–564, doi:10.1016/j.bcp.2005.10.044 (2006).

26 Cumming, R. C. et al. Protein disulfide bond formation in the cytoplasm during oxidative stress. J Biol Chem 279, 21749–21758, doi:10.1074/jbc.M312267200 (2004).

27 Hochgrafe, F. et al. S-cysteinylation is a general mechanism for thiol protection of Bacillus subtilis proteins after oxidative stress. J Biol Chem 282, 25981–25985, doi:10.1074/jbc.C700105200 (2007).

28 Takeuchi, A. et al. A human erythrocyte-derived growth-promoting factor with a wide target cell spectrum: identification as catalase. Cancer Res 55, 1586–1589 (1995).

29 Bolduc, J. A. et al. Novel hyperoxidation resistance motifs in 2-Cys peroxiredoxins. J Biol Chem 293, 11901–11912, doi:10.1074/jbc.RA117.001690 (2018).

30 Haynes, A. C., Qian, J., Reisz, J. A., Furdui, C. M. & Lowther, W. T. Molecular basis for the resistance of human mitochondrial 2-Cys peroxiredoxin 3 to hyperoxidation. J Biol Chem 288, 29714–29723, doi:10.1074/jbc.M113.473470 (2013).

31 Peskin, A. V. et al. Glutathionylation of the Active Site Cysteines of Peroxiredoxin 2 and Recycling by Glutaredoxin. J Biol Chem 291, 3053–3062, doi:10.1074/jbc.M115.692798 (2016).

32 Peskin, A. V. et al. Modifying the resolving cysteine affects the structure and hydrogen peroxide reactivity of peroxiredoxin 2. J Biol Chem 296, 100494, doi:10.1016/j.jbc.2021.100494 (2021).

33 Cavallini, D., De Marco, C. & Dupre, S. Luminol chemiluminescence studies of the oxidation of cysteine and other thiols to disulfides. Arch Biochem Biophys 124, 18–26, doi:10.1016/0003-9861(68)90299-3 (1968).

34 Steinhilper, R., Hoff, G., Heider, J. & Murphy, B. J. Structure of the membrane-bound formate hydrogenlyase complex from Escherichia coli. Nat Commun 13, 5395, doi:10.1038/s41467-022-32831-x (2022).

35 Langford, T. F., Deen, W. M. & Sikes, H. D. A mathematical analysis of Prx2-STAT3 disulfide exchange rate constants for a bimolecular reaction mechanism. Free Radic Biol Med 120, 239–245, doi:10.1016/j.freeradbiomed.2018.03.039 (2018).

36 Honzejkova, K., Kosek, D., Obsilova, V. & Obsil, T. The cryo-EM structure of ASK1 reveals an asymmetric architecture allosterically modulated by TRX1. Elife 13, doi:10.7554/eLife.95199 (2024).

