## supplementary data for "Challenges in reconstituting the peroxiredoxin 2:STAT3 transient redox-relay complex *in vitro*"

**Supplementary Table 1. List of primers used in this study.**

| Purpose | Direction (forward:Fwd; reverse: Rev) | Sequence (5' → 3') |
|---|---|---|
| HiFi DNA assembly of Prdx2 in pD441 | Prdx2-Fwd | cacagagaacagattggtggtatggcctccggtaacgc |
| | Prdx2-Rev | gtgcggccgcctcgagttactaattgtgtttggagaaatattccttgctgtc |
| | pD441-Fwd | gaatatttctccaaacacaattagtaactcgaggcggccgc |
| | pD441-Rev | cgcgcgttaccggaggccataccaccaatctgttctctgtgag |
| HiFi DNA assembly of CF-STAT3 in pD441 | CF-STAT3-Fwd | cacagagaacagattggtggtgtggtgacggagaagcagc |
| | CF-STAT3-Rev | gtgcggccgcctcgagttatggccgacaatactttccgaatg |
| | pD441-Fwd | cggaaagtattgtcggccataactcgaggcggccg |
| | pD441-Rev | gctgcttctccgtcaccacaccaccaatctgttctctgtgagc |

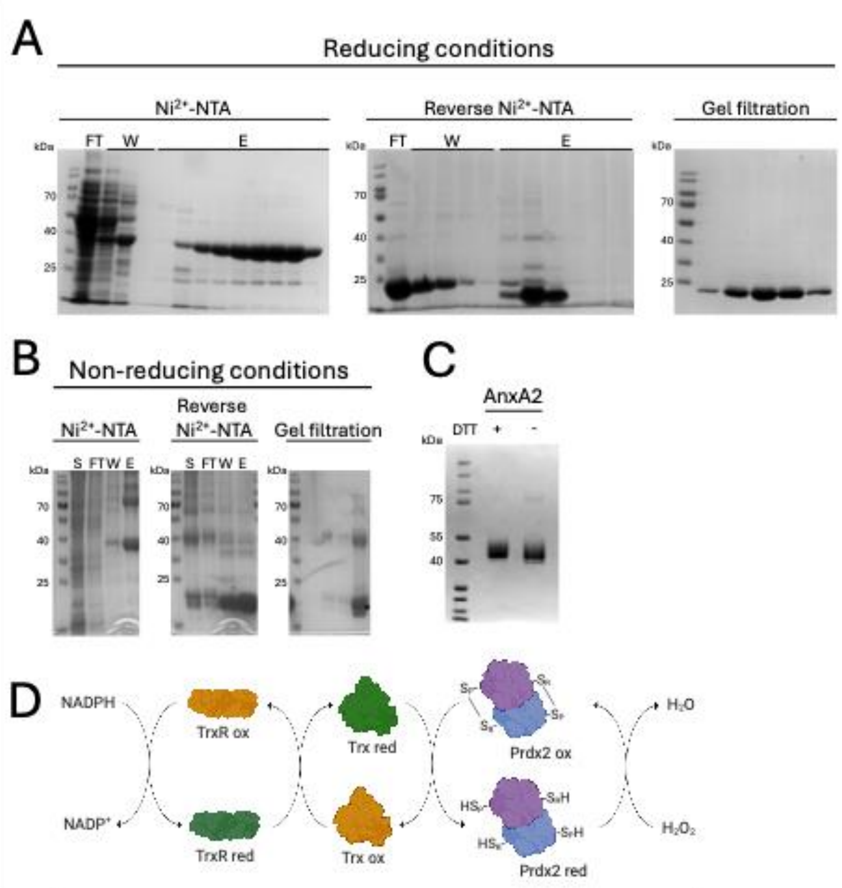


**Supplementary Figure 1. Proteins are successfully purified to homogeneity. (A)** Reducing 12% SDS-PAGE analysis of purified Prdx2 under reducing conditions at each step of the purification process. FT: flow-through; W: wash; E; elution (cropped gel image). **(B)** Non-reducing 12% SDS-PAGE analysis of purified Prdx2 under non-reducing conditions at each step of the purification process. FT: flow-through; W: wash; E; elution (cropped gel image). **(C)** SDS-PAGE analysis of purified Annexin A2 in reducing and non-reducing conditions (cropped gel image). **(D)** Schematic of the NADPH-coupled assay used to assess Prdx2 peroxidase activity.

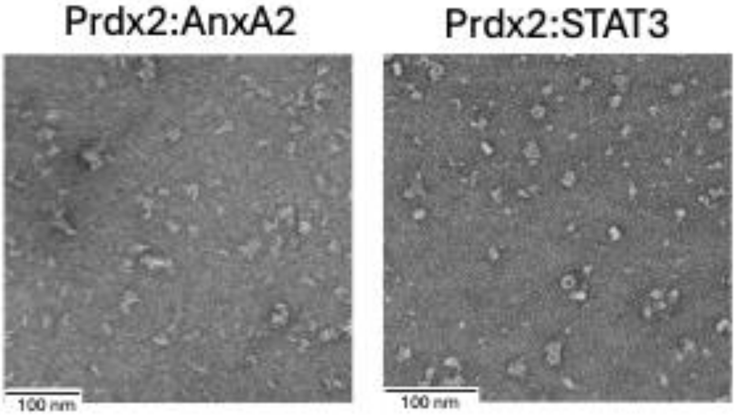


**Supplementary Figure 2. Negative-stain EM images of Prdx2:STAT3 mixtures reveal diffuse, heterogeneous particles with no evidence of higher-order assemblies.** Incubation of Prdx2 and AnxA2 does not promote the formation of discrete ternary complexes. Micrographs show increased aggregation and sample heterogeneity, consistent with non-productive interactions or misfolding. Pre-reduced proteins were mixed in a 2:2 molar ratio in low salt. Micrographs were acquired under reducing conditions at a protein concentration of 0.05 mg/mL.

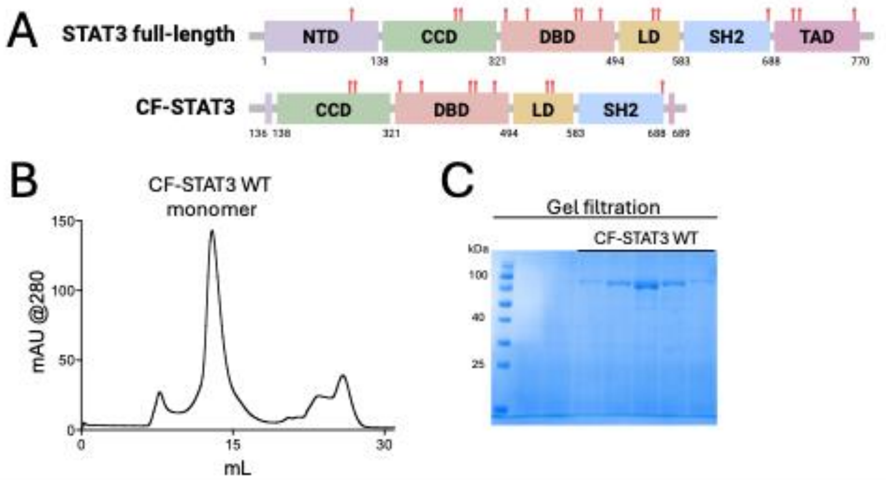


**Supplementary Figure 3. CF-STAT3 does not show degradation products. (A)** Full-length STAT3 and CF-STAT3 (AA 136-689) representation. Domains: helical N-terminal domain (NTD), coiled-coil domain (CCD), central DNA-binding domain (DBD), linker domain (LD), Src homology 2 (SH2) domain, and C-terminal transactivation domain (TAD). Cys residues are shown as red sticks in their positions. Figure created using BioRender. **(B and C)** 12% SDS-PAGE analysis and SEC profiles of CF-STAT3, showing high purity and lack of degradation products. CF-STAT3 remains monomeric and homogeneous across preparations (cropped gel image).

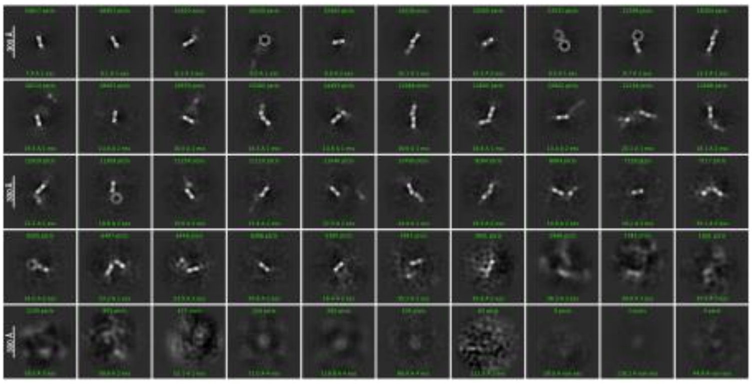

**Supplementary Figure 4. 2D class averages of selected particles show that only Prdx2 particles are present as isolated particles.** Micrographs were acquired under reducing anaerobic conditions at a protein concentration of 0.3 mg/mL. Cryo-EM micrographs were analyzed using CryoSparc v4.6.0.
